# Proficiency of WHO Global Foodborne Infections Network External Quality Assurance System participants in the identification and susceptibility testing of thermo-tolerant *Campylobacter* spp. from 2003-2012

**DOI:** 10.1101/359794

**Authors:** Susanne Karlsmose Pedersen, Jaap A. Wagenaar, Håkan Vigre, Louise Roer, Matthew Mikoleit, Awa Aidara-Kane, Amy L. Cawthorne, Frank M. Aarestrup, Rene S. Hendriksen

## Abstract

*Campylobacter* spp. are food- and water borne pathogens. While rather accurate estimates for these pathogens are available in industrialized countries, a lack of diagnostic capacity in developing countries limits accurate assessments of prevalence in many regions. Proficiency in the identification and susceptibility testing of these organisms is critical for surveillance and control efforts. The aim of the study was to assess performance for identification and susceptibility testing of thermo-tolerant *Campylobacter* among laboratories participating in the World Health Organization (WHO) Global Foodborne Infections Network (GFN) External Quality Assurance System (EQAS) over a nine year period.

Participants (primarily national level laboratories) were encouraged to self-evaluate performance as part of continuous quality improvement.

The ability to correctly identify *Campylobacter* spp. varied by year and ranged from 61.9 % (2008) to 90.7 % (2012), and the ability to correctly perform antimicrobial susceptibility testing (AST) for *Campylobacter* spp. appeared to steadily increase from 91.4 % to 93.6 % in the test period (2009-2012).

Poorest performance (60.0 % correct identification and 86.8 % correct AST results) was observed in African laboratories.

Overall, approximately 10 % of laboratories reported either an incorrect identification or antibiogramme. As most participants were (supra)-national reference laboratories, these data raise significant concerns regarding capacity and proficiency at the local, clinical level. Addressing these diagnostic challenges is critical for both patient level management and broader surveillance and control efforts.

## Introduction

Campylobacteriosis in humans is typically presents as acute diarrhea with fever. However, more significant sequelae such as Guillain-Barré syndrome (GBS), reactive arthritis (ReA) and irritable bowel syndrome (IBS) have been reported following *Campylobacter* gasteroenteritis (4). Most human cases are caused by thermo-tolerant *Campylobacter* spp. which are zoonotic bacteria found in animals such as poultry, cattle and pigs as well as contaminants of various foodstuffs and water (1, 2).

*Campylobacter jejuni* or *Campylobacter coli* are the most commonly implicated species and campylobacteriosis is the most frequently reported bacterial foodborne illness in most developed countries. However, data from developing countries data is often limited by a lack of diagnostic capacity (1, 3, 4).

Antimicrobials are typically not indicated for mild/moderate enteritis in otherwise healthy individuals. However, antimicrobial therapy may be warranted for severe or bloody diarrhea. Antimicrobials are also used in the management of extra-intestinal (invasive) infections. The macrolides (e.g. erythromycin or azithromycin) are commonly used for empiric treatment of campylobacteriosis and fluoroquinolones (e.g. ciprofloxacin) may be a second line therapy for adults. Accurate antimicrobial susceptibility testing is critical for developing empiric therapy guidelines and monitoring emerging resistance. Increasing antimicrobial resistance (AMR), especially multidrug resistance to fluoroquinolones and azithromycin significantly limits treatment options for severe/invasive disease. Access to last line drugs such as carbapenems is often beyond the reach of many in the developing world (5).

Since 2000, the World Health Organization (WHO) Global Foodborne Infections Network (GFN) (formerly WHO Global Salm-Surv (WHO GSS)) has functioned as an international platform to enhance the capacity of countries to detect, control and prevent foodborne and other enteric infections. Part of this capacity building work has focused on identification and susceptibility testing of *Campylobacter*. Since 2000, WHO GFN has offered members the opportunity to participate at no cost in an annual External Quality Assurance System (EQAS). Although the primary focus of the EQAS is serotyping and antimicrobial susceptibility testing (AST) of *Salmonella*; identification and AST of *Campylobacter* is included as a separate module. Laboratories may choose to participate in all or some components. Approximately 200 laboratories participate in one or more components of the EQAS. Of these, approximately 25 % will participate in the *Campylobacter* module. The WHO GFN program focuses activities mainly at reference level facilities (supranational, national, or subnational). While some of these facilities may perform clinical testing; clinical diagnostic laboratories typically do not participate in EQAS (6,7). Participants report results electronically and receive their results immediately. Participants are encouraged to utilize their results as part of continuous quality improvement.

The aim of this paper is to summarise and describe temporal and geographic trends in the performance of the *Campylobacter* component of the EQAS (identification and AST) observed between 2003-2012.

## Materials and Methods

The *Campylobacter* identification component has included *C. jejuni* and *C. coli* since 2003 and *Campylobacter lari* (from 2003-2008). AST of *C. jejuni* and *C. coli* have been included since 2009. Due to the limited availability of epidemiological cut off values (ECOFFs) for other *Campylobacter* spp., the AST component, only includes *C. jejuni* and *C. coli.* Since 2003, the Technical University of Denmark, National Food Institute (DTU Food) in collaboration with members of the WHO GFN steering committee have organized this proficiency test annually (except 2005). DTU Food coordinated the selection of test strains and verified the identification and AST of test strains. Results obtained by DTU-Food were reconfirmed in a blinded manner by a referee laboratory (United States’ Centers for Disease Control and Prevention (CDC)). Further details on the preparatory work and the EQAS-setup are described in the annual EQAS reports available on the Internet (http://www.who.int/gfn/activities/eqas/en/).

While the target audience for the EQAS is national public health, food and veterinary reference laboratories, in special instances (particularly in countries without a designated referral laboratory) the organizers occasionally permit a select number of clinical and/or research laboratories to participate in the EQAS. EQAS participants have the option to participate in all or some of the components. Participants in the WHO GFN proficiency test for identification and/or AST of *Campylobacter* receive two vials each containing a lyophilized *Campylobacter* isolate (challenge strains). In addition, all laboratories participating in the *Campylobacter* AST component were provided an isolate of *C. jejuni* ATCC 33560 upon request. Protocols available on the Internet (http://www.antimicrobialresistance.dk/233-169-215-eqas.htm) described how to revive and test the isolates and referred to a manual on sub-culturing and maintenance of quality control (QC) strains.

For the identification component, the protocol specified that the laboratory’s routine methods should be applied. Laboratories were free to utilize conventional phenotypic identification, molecular identification, or a combination of methods for identification. Laboratories who participated only in the AST component (did not perform identification), were provided upon request the identification of the coded *Campylobacter* strains.

While multiple methods were used for identification; during this period, time validated methods for susceptibility testing of *Campylobacter* by disk diffusion or E-test were not internationally available. As such only broth or agar dilution methods (MIC) were accepted for the AST component. Epidemiological cut-off values (ECOFFs) for the interpretation of disk diffusion zones for ciprofloxacin, erythromycin and tetracycline against *C. coli* as well as ciprofloxacin, erythromycin, tetracycline, and gentamycin for *C. jejuni* are now incorporated into EUCAST guidelines Participants could test and submit results for chloramphenicol (CHL), ciprofloxacin (CIP), erythromycin (ERY), gentamicin (GEN), nalidixic acid (NAL), streptomycin (STR), and tetracycline (TET). The protocol listed the interpretative criteria applied for this EQAS, i.e. ECOFFs according to EUCAST (http://www.eucast.org) which allowed two categories of characterization (resistant [non-wildtype], R or susceptible [wildtype], S) for the *C. jejuni* and *C. coli* test strains.

The WHO GFN EQAS was set-up as a self-evaluating system in which participants directly upon submission of results received a report comparing their obtained results to quality assured and verified expected results and itemizing the laboratory’s eventual deviations. Deviations for the identification component were reported as incorrect results and no attempt was made to quantify their severity. For the AST component, the acceptance limit was set at 5 % deviations, i.e. one deviation would categorize the laboratory’s results as unacceptable. The analysis was based on assigning all results the same level of influence, i.e. disregarding the impact of the variation in the selection of test strains from year to year (the susceptibility testing of some strains could be more difficult than others) and the varying participation levels (some years, more laboratories participated compared to other years).

For each of the world regions, the annual proportion of correctly identified species were analyzed for i) significant variation between the years 2003 to 2012 using the function Fisher.test in R and ii) a time-trend from 2003 to 2012 by performing a chi-square test for trend in proportions using the function prop.trend.test from the R-package stats. Next, for each region, the data was aggregated to the overall proportion of correctly identified *Campylobacter* species over the whole period from 2003 to 2012. These data were used to test if the proportion of correctly identified species was different between the regions using the prop.test function in the R-package stats.

To assess potential differences between regions in the AST performance, the proportion of correct AST result for each antibiotic (CHL, CIP, ERY, GEN, NAL, STR and TET) was analyzed for significant differences between regions using the prop.test function in the stat R-package stats.

All presented 95 % confidence intervals for the proportions were obtained using the function binconf in the R-package Hmisc.

## Results

In total, laboratories from 96 countries (Figure 1) participated in the *Campylobacter* identification and/or AST component of the EQAS in one or more iterations from 2003 to 2012 and included national and other reference laboratories from the veterinary, food and public health sector. For the identification component of the EQAS, the number of countries participating each year were: 53 (2003), 62 (2004), 59 (2006), 59 (2007), 63 (2008), 54 (2009), 62 (2010), 57 (2011) and 69 (2012). In some cases multiple laboratories from a country participated in the EQAS (e.g. MoH and MoA laboratories). The cumulative number of laboratories participating in the *Campylobacter* module was: 97, 111, 100, 104, 112, 92, 100, 82 and 112 participants each of the years, respectively.

**Figure 1:**
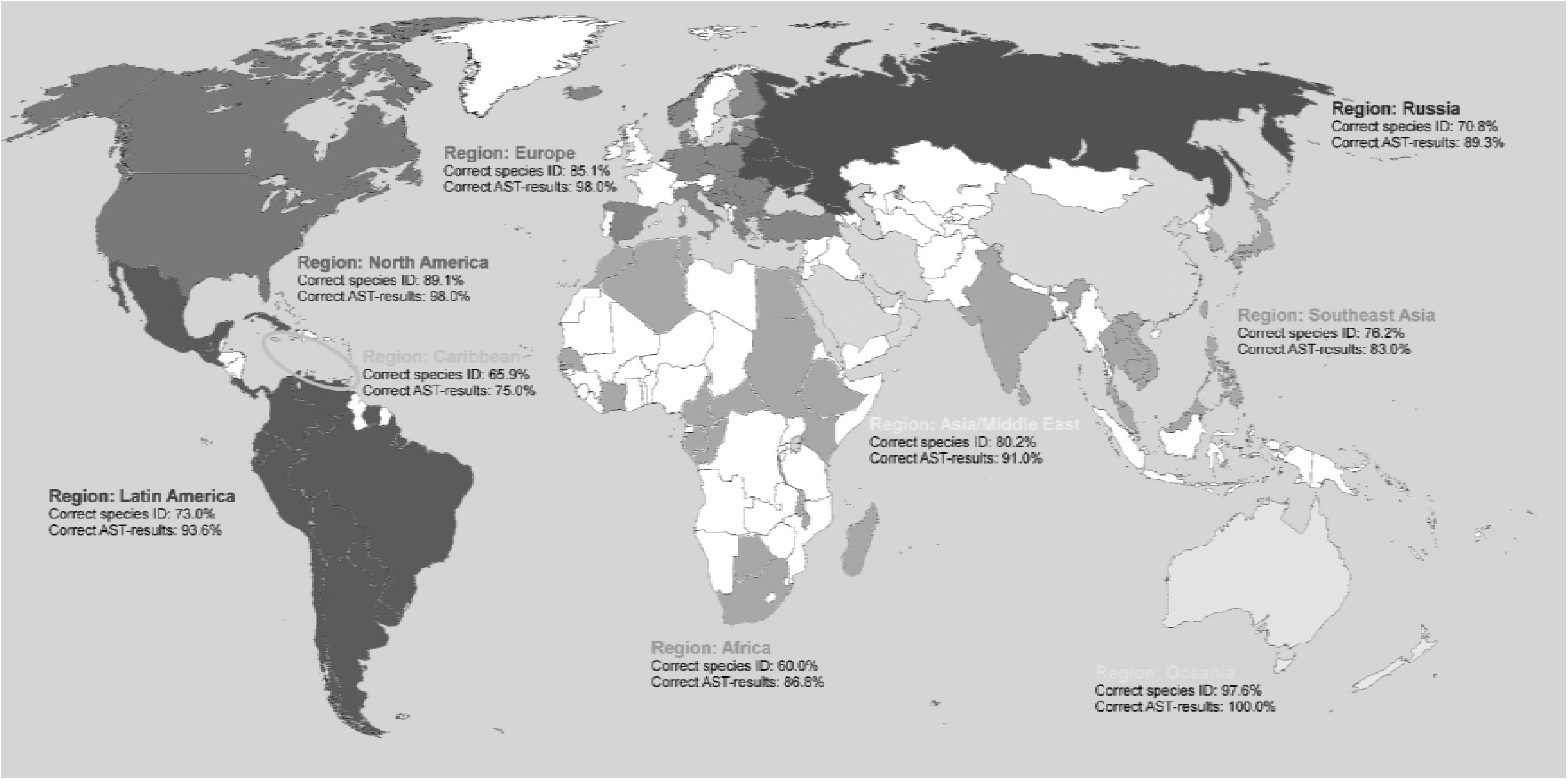
Map indicating the participating countries colored with respect to the region they belong to (Africa, Asia/Middle East, Caribbean, Europe, Latin America, North America, Oceania, Russia or Southeast Asia). Performance with regard to the species identification and antimicrobial susceptibility testing is indicated as average of correct results (%). The 96 participating countries included the following: Africa (Algeria, Botswana, Cameroon, Central African Republic, Congo Rep. of, Egypt, Ethiopia, Gambia, Gabon, Ivory Coast, Kenya, Madagascar, Malawi, Mauritius, Morocco, Senegal, South Africa, Sudan, Tunisia); Asia & Middle East (China, Iran Islamic Rep. of, Israel, Kuwait, Oman, Saudi Arabia); Caribbean (Barbados, Grenada, Jamaica, Trinidad and Tobago); Europe (Bosnia and Herzegovina, Bulgaria, Croatia, Cyprus, Czech Republic, Denmark, Estonia, Finland, Germany, Greece, Hungary, Iceland, Italy, Latvia, Lithuania, Luxembourg, Macedonia, Malta, Rep. of Moldova, Netherlands, Norway, Poland, Romania, Serbia, Slovakia, Slovenia, Spain, Turkey); Latin America (Argentina, Bolivia, Brazil, Chile, Colombia, Costa Rica, Cuba, Ecuador, El Salvador, Guatemala, Mexico, Panama, Paraguay, Peru, Suriname, Uruguay, Venezuela); North America (Canada, United States of America); Oceania (Australia, New Caledonia, New Zealand); Russian region (Belarus, Georgia, Russian Federation, Ukraine); Southeast Asia (Brunei Darussalam, Cambodia, India, Japan, Korea Rep. of, Lao Dem. Rep. of, Malaysia, Philippines, Singapore, Sri Lanka, Taiwan, Thailand, Viet Nam).

For the AST of *Campylobacter* in the years 2009, 2010, 2011 and 2012, the numbers of participating countries from each region were: Africa (2, 2, 7, 4); Asia & Middle East (1, 1, 1, 3); Caribbean (0, 0, 0, 1); Europe (7, 9, 7, 11); Latin America (4, 7, 6, 6); North America (1, 2, 2, 2); Oceania (0, 0, 1, 0); Russian region (0, 1, 1, 0); Southeast Asia (4, 5, 4, 6). The cumulative number or laboratories participating from each country was: 25, 37, 38 and 47 participants in total each of the years, respectively. In all, 18 laboratories participated twice, 14 participated three times and 11 participants took part in all four *Campylobacter* AST iterations from 2009 to 2012.

The overall ability to correctly identify *Campylobacter* spp. fluctuated over the years 2003 to 2012 (Figure 2), and when focusing at *C. coli* and *C. jejuni* between the regions, a significant difference could be identified in Europe and Latin America in the proportion of correctly identified *Campylobacter* species between years (data not shown). No time-related trend was identified in Europe nor Latin America, and other factors (such as new laboratories joining or previous participants leaving the program) likely contributed to this variation. Significant variation between the years was not found in any other regions.

**Figure 2:**
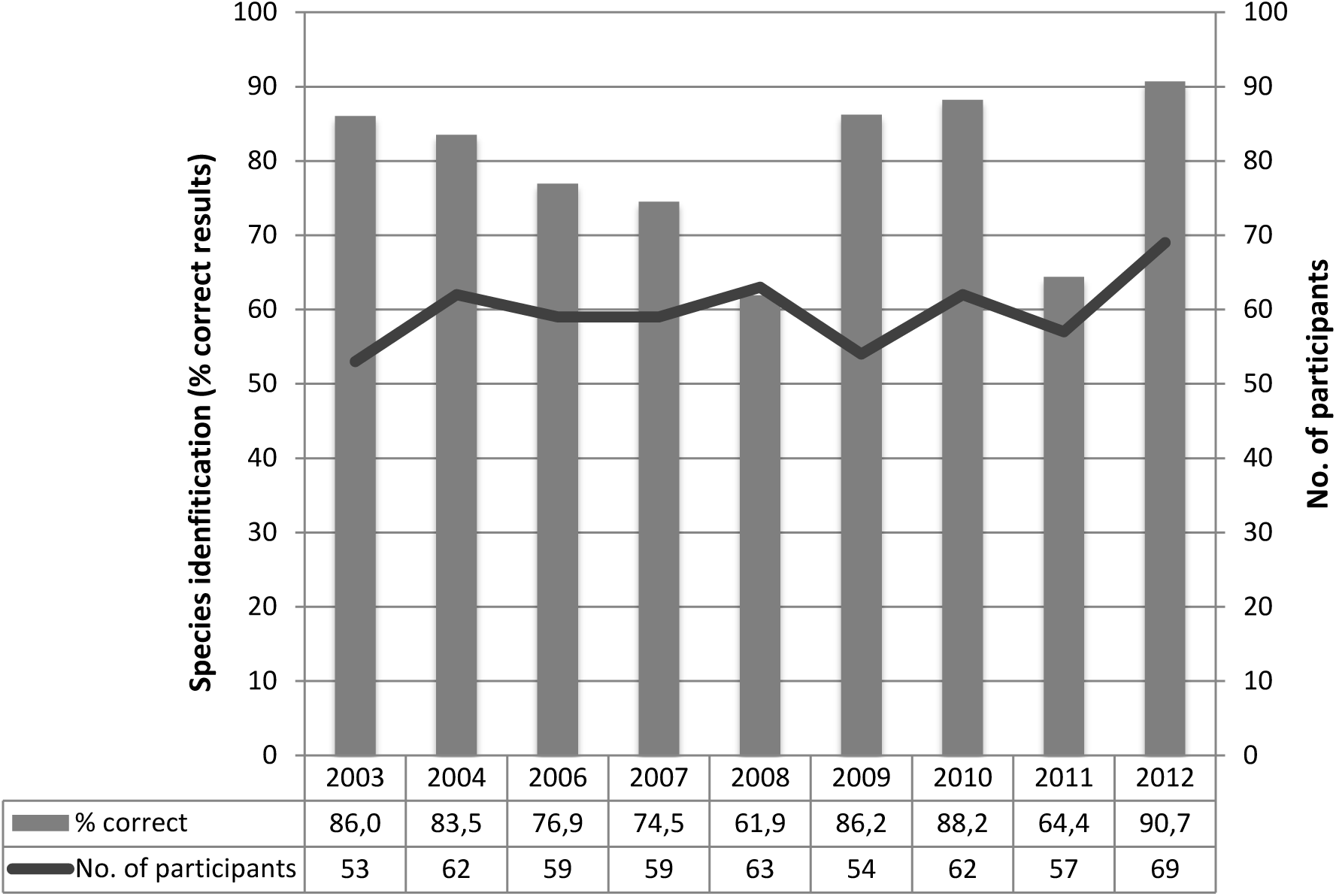
Species identification of *Campylobacter*; summary of the performance per year of results covering all nine participating regions (Africa, Asia/Middle East, Caribbean, Europe, Latin America, North America, Oceania, Russia, Southeast Asia).

There was a strong significant difference between the regions as to the proportion of correctly identified *Campylobacter* species, with Oceania and North America exhibiting the highest, and Africa and the Caribbean the lowest proportions of correctly identified *Campylobacter* species (Figure 3). It is important to consider that in some regions (e.g. Oceania), participation was limited (n=1) and may fail to truly reflect regional capacity.

**Figure 3:**
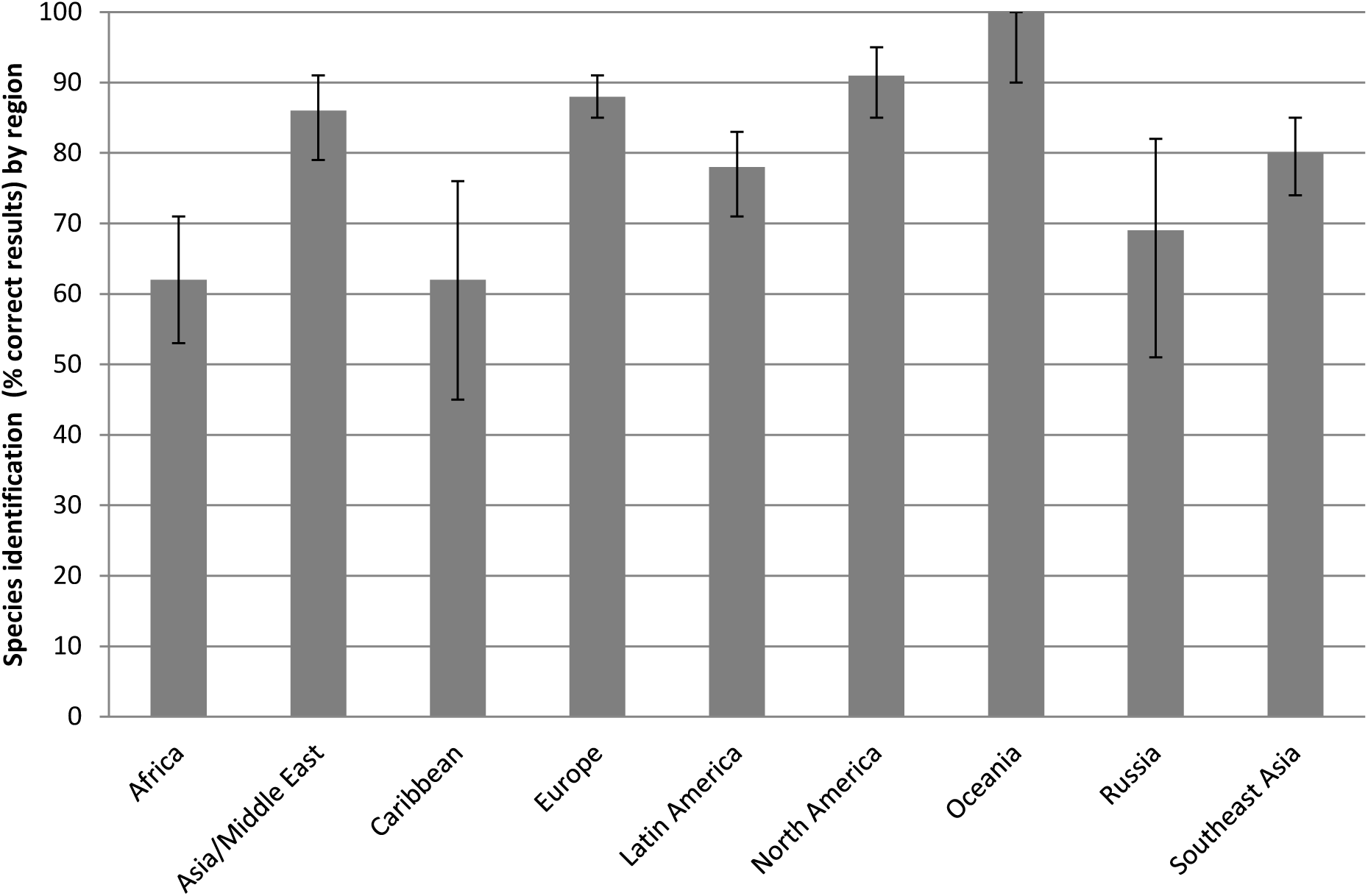
Species identification of *Campylobacter coli* and *C. jejuni;* summary of the performance per region of results over the years 2003-2012 (excl. 2005).

In the iterations from 2003 to 2012, identifications were correctly performed between 59.3 % (2011; *C. coli;* N = 81) and 96.4 % (2012; *C. jejuni;* N = 112) with an average over the years of 79.3 % (based on the result of two isolates per year, i.e. 18 isolates in total, of *C. jejuni, C. coli* and *C. lari*; total number of observations, N = 1,736). Comparing between the species, it appears that participants are able to correctly identify *C. jejuni,* (90.4 % correct) than *C. coli* and *C. lari* (74.1 % and 68.8 % correct identifications, respectively). This result is not unexpected as typical *C. jejuni* is readily identified by hippurate hydrolysis whereas other species require additional, more complex tests.

Subsequent to the validation of the submitted data, 1,565 AST results could be included for analysis for the eight *Campylobacter* isolates included in the four iterations from 2009 to 2012. Of these, 7.3 % deviated from the expected. For all antimicrobials included in the AST performance test, there was a significant difference between the performance of the regions, where Europe exhibited the highest correspondence with the expected results and Southeast Asia the lowest (Figure 4).

**Figure 4:**
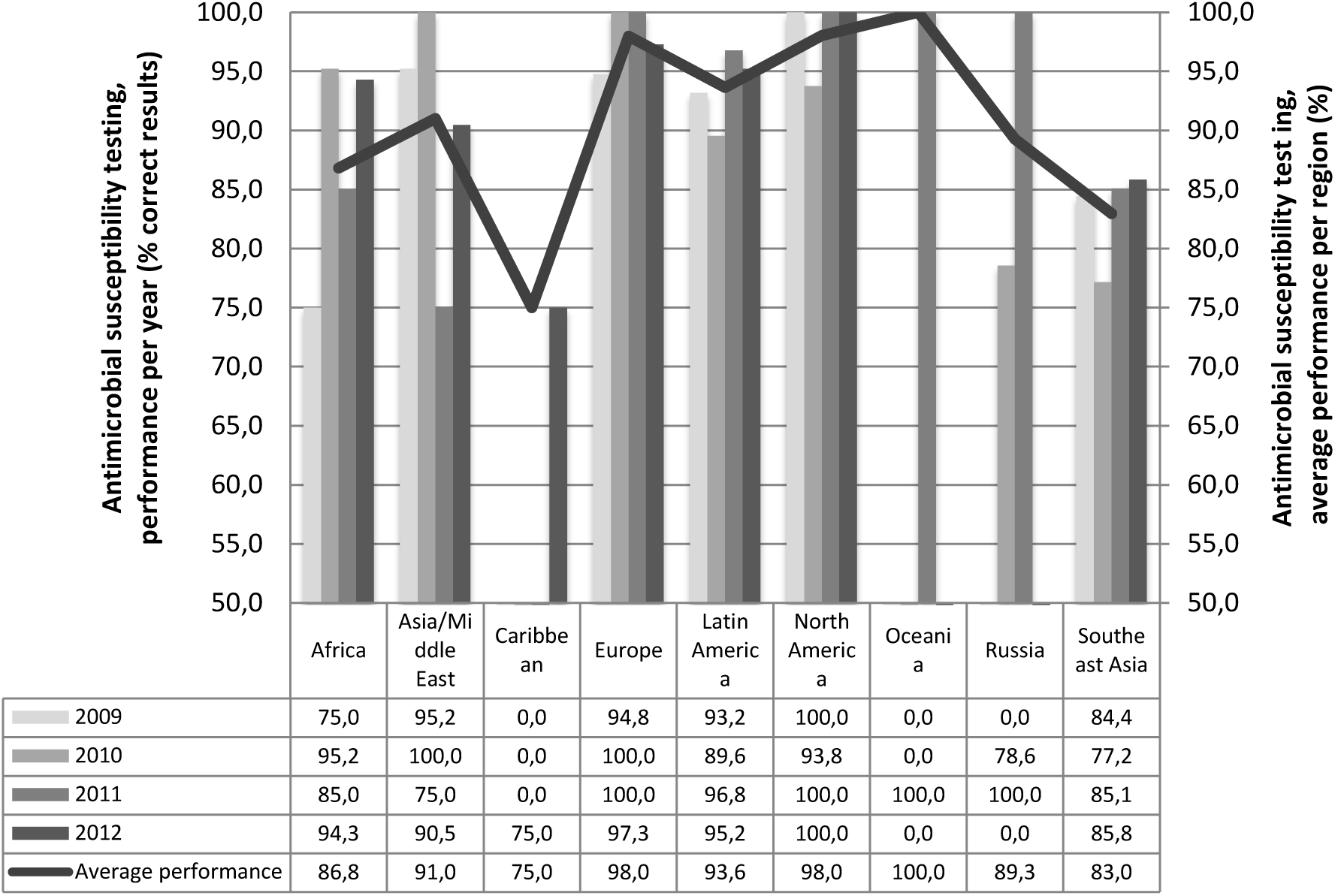
Antimicrobial susceptibility testing of *Campylobacter*; the performance per year and region.

Deviations appeared to be in particular caused by streptomycin with a deviation level at 11.5 %, but also for erythromycin, tetracycline and ciprofloxacin, fairly high deviation levels were seen (8.5 %, 8.1 %, and 6.4 % respectively). In particular, one bacterial strain caused a high level of deviations, i.e. WHO 2009 C-9.2 (resistant to CIP, NAL, and TET) for which 11.3 % of the submitted results deviated from the expected. A low deviation level (4.2 %) was obtained for tests towards chloramphenicol, towards which all the test strains were susceptible. A comparison of the obtained results from all laboratories which participated in the AST component in one or more of the four iterations, indicated a slight increase in performance (Figure 5). Disregarding results from the Caribbean and Oceania due to the limited number of submitted results, the summary of the four years’ results per region provided an indication of a generally low performance of the participants in the Southeast Asian region and the African region, with levels at 17.0 % and 13.2 % deviations, respectively. In 2012, however, all regions except Southeast Asia (deviation level at 14.2 %) exhibited deviation levels lower than 10 % (the Russian region did not participate in the 2012 iteration).

**Figure 5:**
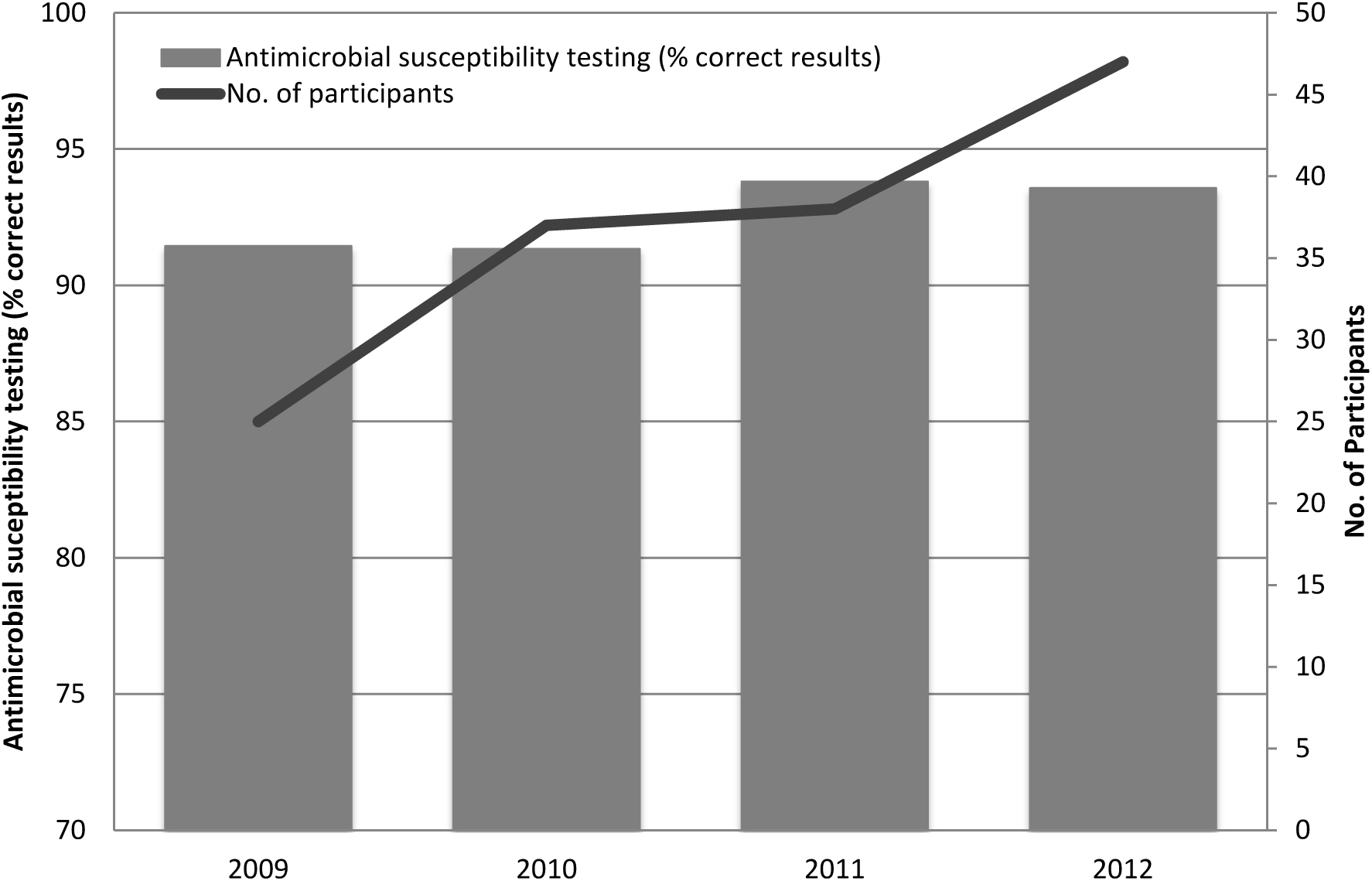
Antimicrobial susceptibility testing of *Campylobacter*; summary of the performance per year of results covering all nine participating regions (Africa, Asia/Middle East, Caribbean, Europe, Latin America, North America, Oceania, Russia, Southeast Asia).

In the four iterations, 51 (75 %) laboratories uploaded one or more values for the QC reference strain, *C. jejuni* ATCC 33560 suggesting that 25 % of the labs did not test QC strain for reference. Of the submitted values for the QC reference strain, an average of 17.2 % were out of the QC range when evaluated towards one of the validated methods described by CLSI (e.g. VET01-A4) (8). Analysis of regional differences in this context reveals Africa as the region with the highest level of laboratories submitting AST results for the *Campylobacter* test strains without submitting results for the *C. jejuni* ATCC 33560 reference strain.

## Discussion

The proficiency test results are intended to be utilized for continuous quality improvement. However, some participating laboratories did not demonstrate improvement in identification or AST over time. Information about corrective actions implemented in the individual laboratories based on the deviations in their evaluation report could have added to the analysis but was unfortunately not available. This proficiency test shows a worrisome tendency where even national reference laboratories in regions, normally anticipated to perform flawless, have approximately 10 % incorrect results in both tests. Given the complex microbiology of *Campylobacteriaceae* and the fastidious nature of these organisms, when these results are extrapolated to lower level facilities such as clinical laboratories, the ability to correctly identify and antimicrobial susceptibility test *Campylobacter* is likely significantly higher than the 10 % error observed among participating laboratories. In addition, *Campylobacter* results were only received from ~25 % of the total number of participating laboratories in the WHO GFN EQAS. This suggests that nearly 75 % of participants lack basic capacity for testing *Campylobacter.* These findings are worrying in light of the importance of *Campylobacter* as a foodborne pathogen. Thus, actions are still required to improve the performance of laboratories and report correct data on *Campylobacter* worldwide.

Performance (pass/fail) criteria were not specified for the *Campylobacter* identification module, though the average of 79.3 % correctly identified strains indicates that some laboratories would need to assess their routine to improve their performance. Assessing the general performance of the AST component of all participating laboratories over the four years, the average deviation level at 7.3 % exceeds the defined acceptance limit of 5 %. The development in the annual deviation levels; however, could not indicate a trend with statistical significance. For the AST, the high level of incorrectly reported results is critical, especially for macrolides (8.5 %) and fluoroquinolones (6.4 %), since these two antimicrobials classes are the preferred choice for treatment.

The frequent incidence of *Campylobacter* diarrhea, increasing drug resistance, and the potential for long term sequelae, highlight the importance of accurately understanding the socio-economic burden of campylobacteriosis (4). Increased competence of reference laboratories for identification and AST *C. jejuni* and *C. coli* therefore support disease surveillance and control programs. However, the ability of a referral center to impact surveillance is contingent upon an isolate/specimen flow from lower level laboratories. On average, only 25 % of EQAS participants reported results for *Campylobacter,* suggesting widespread gaps in capacity. The advancement in whole genome sequencing and *in silico* bioinformatics tools combined with lower prices, is a promising development in enhancing the ability of national reference laboratories to correctly identify and susceptibility testing *Campylobacter* in the future. Similarly, molecular assays and other culture independent tests may increase surveillance capacity at the clinical level. While these technologies currently are not a substitute for culture, they may provide estimates of burden and help determine which specimens should be subjected to conventional culture.

The self-evaluation design of this proficiency test was intended to challenge the participating laboratories to assess their current identification and AST methods for *Campylobacter* also allowing them to include the proficiency test outcome as an external quality assessment of the relevant methods. Self-evaluation is a concept well-known to laboratories following a quality assurance standard requiring quality control procedures e.g. ISO/IEC 17025 (9) and might include monitoring the validity of test results by regular use of internal quality control using reference materials or participation in proficiency-testing programmes. Apart from the obvious connection to the WHO GFN EQAS as a proficiency test, laboratories participating in the programme are offered material for internal control in the form of both the certified ATCC reference strain, *C. jejuni* ATCC 33560, and the test strains which can be regarded as internal control strains and consequently should be stored and maintained.

In addition to self-evaluation, the possibility of introducing approaches like mentoring of participants, training courses, and E-learning could be explored and suggested to the participants. The question of resources must, however, be considered, for example, mentoring of participants appears to be a rewarding but also is a resource demanding approach of capacity building Regional follow-up is likely to be a rewarding approach and should be based on evaluation of regional needs and challenges. Especially for the African region and Southeast Asia, it appears that specific follow-up is required. For example, the submitted results for the AST component indicate that many laboratories did not perform adequate internal quality control (17.2 % of submitted results for the ATCC reference strain were out of range), which is why WHO GFN capacity building efforts focus at encouraging the maintenance of relevant quality assurance as part of the laboratory routines. Internal laboratory QA ensures minimization of variable factors influencing the obtained result for a test strain. These factors include the media content, the activity of the antimicrobial, and the testing of a QC reference strain according to an internationally recognized standard (e.g. CLSI). Laboratories that have introduced relevant quality assurance of the variable factors facilitate a good performance. For all laboratories performing AST of *Campylobacter* species, testing of the *C. jejuni* reference strain (ATCC33560) should be a routine QA measure providing quality control for both the method and the reagents. Moreover, results outside the quality control ranges should always induce appropriate follow-up.

In conclusion, this annually provided proficiency test supports the identification and AST of *Campylobacter* and allows national reference laboratories free of charge to evaluate their obtained results by comparison to quality assured and verified expected results. Overall, we found that global ability to correctly identify *Campylobacter* spp. fluctuated over the years up to 90.7 % and the ability to correctly perform AST appeared to steadily increase to 93.6 %. African laboratories followed by Southeast Asian had the lowest performance in both identifying the *Campylobacter* spp. conducting AST. Our results reveal a worrisome tendency where approximately 10 % of laboratories report either an incorrect diagnosis or antimicrobial susceptibility profile for treatment. This will compromise the ability to correctly diagnose illness, effectively treat patients and will provide unreliable data for pathogen and AST surveillance systems if not attended.

## Acknowledgement

This work was funded by the World Health Organization Global Foodborne Infections Network (http://www.who.int/gfn) and by COMPARE, which has received funding from the European Union’s Horizon 2020 research and innovation programme under grant agreement No 643476. Thanks to Arne B. Jensen for setting up the submission database for capturing and evaluating the results of the EQAS participants.

